# Overcoming leak sensitivity in CRISPRi circuits using antisense RNA sequestration and regulatory feedback

**DOI:** 10.1101/2022.03.24.485671

**Authors:** David A. Specht, Louis B. Cortes, Guillaume Lambert

## Abstract

The controlled binding of the catalytically-dead CRISPR nuclease (dCas) to DNA can be used to create complex, programmable transcriptional genetic circuits, a fundamental goal of synthetic biology. This approach, called CRISPR interference (CRISPRi), is advantageous over existing methods because the programmable nature of CRISPR proteins enables the simultaneous regulation of many different targets without crosstalk. However, such gene circuit elements are limited by 1) the sensitivity to leaky repression of CRISPRi logic gates and 2) retroactive effects owing to a shared pool of dCas proteins. By utilizing antisense RNAs (asRNAs) to sequester guide RNA transcripts, as well as CRISPRi feedback to self-regulate asRNA production, we demonstrate a mechanism that suppresses unwanted CRISPRi repression and improve logical gene circuit function in *E. coli*. This improvement is particularly pronounced during stationary expression when CRISPRi circuits do not achieve the expected regulatory dynamics. Further, the use of dual CRISPRi/asRNA inverters restores logical performance of layered circuits such as a double inverter. By studying circuit induction at the single cell level in microfluidic channels, we provide insight into the dynamics of antisense sequestration of gRNA and regulatory feedback on dCas-based repression and derepression. These results demonstrate how CRISPRi inverters can be improved for use in more complex genetic circuitry without sacrificing the programmability and orthogonality of dCas proteins.

## Introduction

A fundamental goal of biological engineering is the implementation of entirely new transcriptional regulatory interactions and gene networks inside a cell. Controlling such networks allows us to endow microorganisms with original engineered behaviors, such as oscillators,^1^ memory elements,^2^ and complex logic functions. ^3^ Despite advances in the standardization of various genetic components in bacteria, including molecular sensors^4^ and terminators,^5^ we still lack standard universal transcriptional processing components which can be reprogrammed to interact with arbitrary regulatory components or be reused within large scale synthetic gene networks.

It has been shown that synthetic transcription factors based on CRISPR interference (CRISPRi)^6–10^ can be used to reprogram cellular function. A catalytically-dead CRISPR protein, designated dCas, can be utilized in bacteria as a programmable transcriptional repressor by blocking RNA polymerase transcription at the CRISPR binding site (Fig. 1A). Using CRISPR as a regulatory element is advantageous because repression can be targeted to any arbitrary DNA sequence as long as a PAM site (protospacer adjacent motif) is present. Further, dCas proteins can be combined with other components to create CRISPR activators (CRISPRa)^11,12^ in addition to repressors. Due to their practically infinite potential for programmability and orthogonality (simultaneous expression without crosstalk), gene circuits using CRISPRi stand to drastically expand the capabilities of synthetic gene networks.

**Figure 1:**
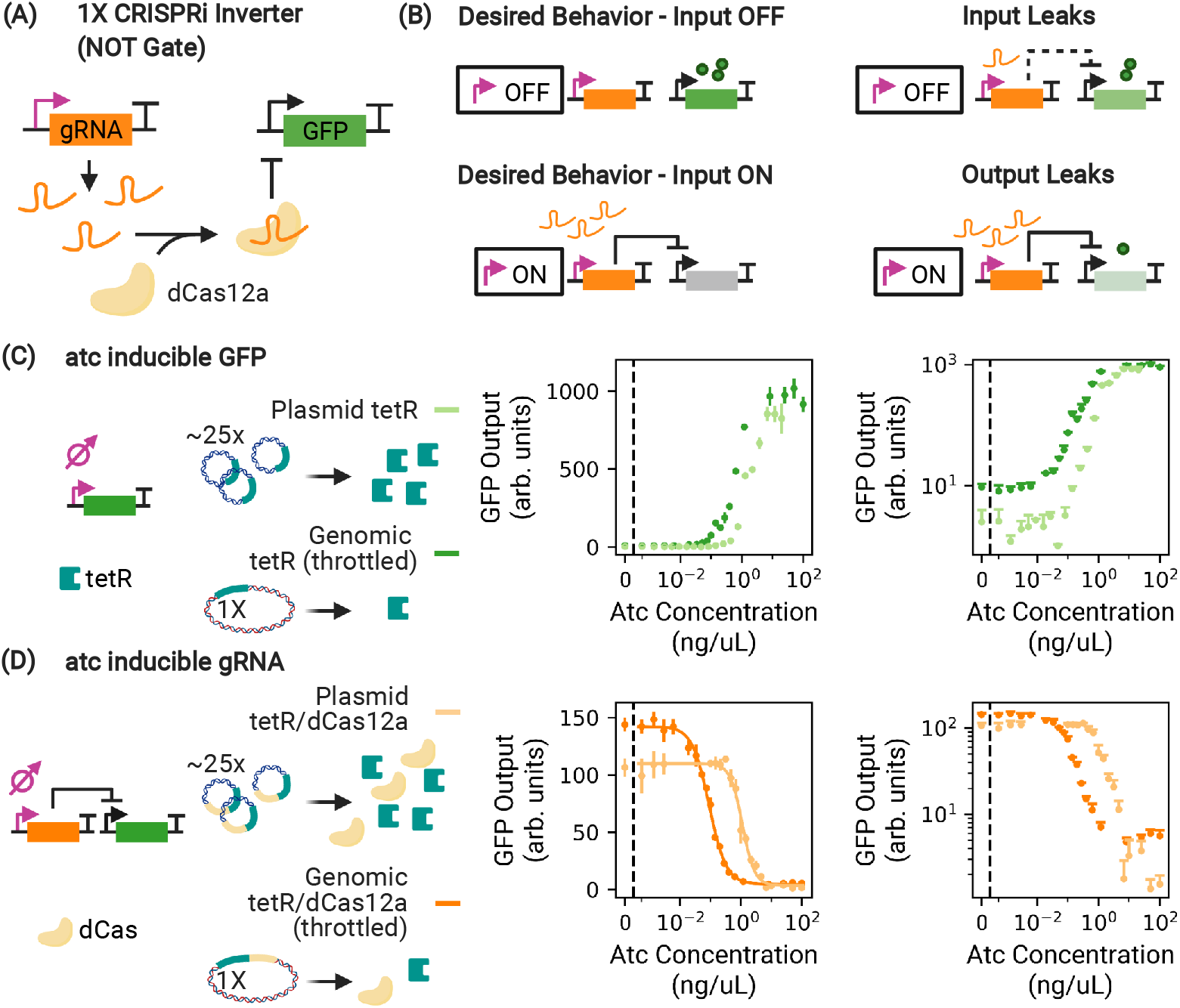
Antisense sequestration as a means to mitigate input leak in dCas-based transcriptional circuits. (A) In its most basic implementation, a CRISPRi-based NOT gate functions by driving the production of a gRNA, which programs dCas to bind to and repress expression from the target promoter. (B) If the input module is an inducible sensor, any basal expression allows the unwanted production of a few gRNA which can efficiently repress the output (Input Leak). Downstream applications can be hindered by incomplete repression by dCas (Output Leak). (C) We throttle tetR availability by expressing it in the genome (dark green), which causes leaky pTet expression at low atc concentration when compared to 20 to 30-fold plasmid (p15A origin) expression (light green). (D) We throttle both tetR and dCas availability, now for the 1X inverter. Throttling dCas decreases sensitivity to leaked gRNA at low atc concentration, increasing the overall dynamic range (dark orange) with respect to high copy plasmid expression of dCas (light orange). However, this decreases the absolute off level of GFP expression, as evident in logspace. Throttling the availability of tetR and dCas increases input and output leak, facilitating study of how these impacts can be mitigated. Curves depicted in (D) and (E) are taken during exponential growth. In linear space displayed error bars are ±1 standard deviation from threefold biological replicates.

However, despite successes in creating dCas-based endpoint^3,13,14^ and dynamic^15–17^ circuits, challenges remain when circuits are scaled up from just a few CRISPRi elements.^18^ In fact, CRISPR’s reprogrammability is also the source of its greatest weaknesses: because a shared pool of dCas proteins is drawn on simultaneously from all active elements, or nodes, in a circuit, it is possible for downstream nodes to interfere with the regulatory activity of those further upstream, an effect called retroactivity.^19–21^ Even if we neglect the effects of retroactivity entirely, CRISPRi circuits are extremely sensitive to transcription leaks^22^ because they lack the nonlinear cooperative response which is necessary to mitigate the impact of leaky repression. We quantify “input leak” as the amount of transcripts expressed when the input promoter is in the off state (i.e. when no atc is present for an inducible pTet promoter). These leaked gRNAs are processed by dCas12a and may bind to their target, reducing GFP expression from the expected maximum (Fig. 1B). Alternatively, “output leak” is the instance where the node processing module (dCas12a + gRNA) ineffectively represses the output, increasing output expression above its expected minimum even at full gRNA induction (Fig. 1B).

In this work, we use 1X CRISPRi inverters derived from *F. novicida* Cas12a^8,9,23^ to demonstrate that CRISPRi in combination with antisense sequestration reduces the impact of retroactivity and the sensitivity to transcription leaks. The benefits are particularly pronounced during post-exponential growth and stationary expression, when accumulation of dCas with leaked gRNA transcripts in a simple inverter cripples circuit performance. We further show that CRISPR’s unique mechanism of action can be used to regulate its own antisense regulation, yielding a regulatory feedback mechanism which further increases dynamic range. Extending this to two inverters connected in series (i.e. a 2X inverter), we also show that antisense sequestration drastically improves the dynamic range of a layered genetic circuit. Lastly, we use microfluidics to study the behavior of these inverters at the single cell level, enabling us to observe how sequestration affects the long equilibration time of CRISPRi circuits. Our results show that antisense sequestration can be used to reduce the impact of leakiness in dCas-based nodes without sacrificing the programmability or orthogonality of CRISPRi synthetic gene circuits.

## Results

### Measuring input and output leak of a 1X CRISPRi inverter

A single dCas CRISPRi inverter functions by driving the production of a gRNA which programs dCas to bind to and suppress expression from a targeted output promoter. ^9^ In our work, the inverter drives GFP production (Fig. 1A). Since the nuclease-dead Cas12a (dCas12a) is expressed constitutively, the output is manipulated via the induction of gRNA transcription using an anhydrotetracycline (atc)-inducible pTet promoter. Thus, in this model system, cells turn from green to white when atc is added and GFP production ceases.

To better understand ways to mitigate input and output leak, we designed a single CRISPRi inverter which purposely suffers from increased leakiness. We achieved this by throttling it twice. First, we reduced tetR availability, which increases input leak and gRNA transcript levels when atc concentrations are low. We compared the performance of pTet-driven GFP expression against a system where tetR is expressed by the same promoter but at 20 to 30-fold higher copy number using a plasmid (p15A origin, (Fig. 1C).^24^ This results in significantly weakened repression at low atc levels due to limited tetR availability which prevents total suppression of GFP production (Fig. 1C). Next, we reduced dCas12a availability, which increases mRNA production at all atc levels (Fig. 1D), by moving dCas12a from the medium-copy plasmid to the genome. This throttles output such that there is imperfect repression at high levels of atc (Fig. 1D). Thus, the performance of 1X inverter is limited both by tetR availability at low atc induction and dCas12a availability at high atc concentration.

Interestingly, because 1X performance is limited by unwanted dCas12a repression in the zero atc state, limiting dCas12a availability increases the absolute dynamic range of the 1X inverter. This is similar to the effect observed in Ref. 21, where the presence of a competitor CRISPRi module increases the dynamic range of a basic inverter by utilizing available dCas12a space when gRNA production is low. Because of this, all circuits in this work are throttled with low genomic dCas12a and tetR availability unless otherwise noted, but the circuit itself is expressed on a plasmid with low, stringently-controlled copy number (pSC101).

### Sequestration of gRNA transcripts reduces input leak sensitivity during stationary expression

To decrease the impact of input leak, we used a format inspired by Ref. 25 and antisense RNA (asRNA) design rules from Ref. 26 to create a hybrid CRISPRi/asRNA system which pairs each CRISPRi node with an asRNA node specifically designed to orthogonally sequester the corresponding gRNA (Figs. 2A, S1). The gRNA is designed to target the −10 site of the target promoter which we know effectively represses transcription based on our previous work.^9^ The asRNA includes a tag which recruits Hfq, a protein which facilitates RNA-RNA interactions.^27^ While previous authors de-repressed dCas9 by binding to a linker located between the sgRNA hairpin and the terminator,^25^ we sequester dCas12a-based gRNAs by binding to a longer sequence which occludes the 20 bp spacer, a 40 bp unique tag, and a portion of the CRISPR repeat sequence. A comparison of occluding different lengths of the gRNA is included in Fig. 2C. Efficacy of sequestration depends on disruption of the repeat hairpin, but this comes at the cost of orthogonal sequestration of unique gRNAs if too many nucleotides of the shared repeat sequence are occluded. Ultimately, we chose to occlude only 9 base pairs of the repeat sequence (Partial Repeat Occlusion, in Fig. 2C), in order to minimize the likelihood of nonorthogonal interactions between asRNAs and noncognate gRNA. We also designed and tested a system to sequester mRNA output, in parallel to gRNA sequestration, which is discussed further in the supplementary text. This system takes advantage of Cas12a’s ability to process its own gRNA, which is an important advantage over Cas9. We take great care to preserve the order and contextual arrangement of nodes. We use sets of 3-character codes in order to specify nodes and their ordering (Supplementary Fig. S1), discussed further in Supplementary Text 1.2.

**Figure 2:**
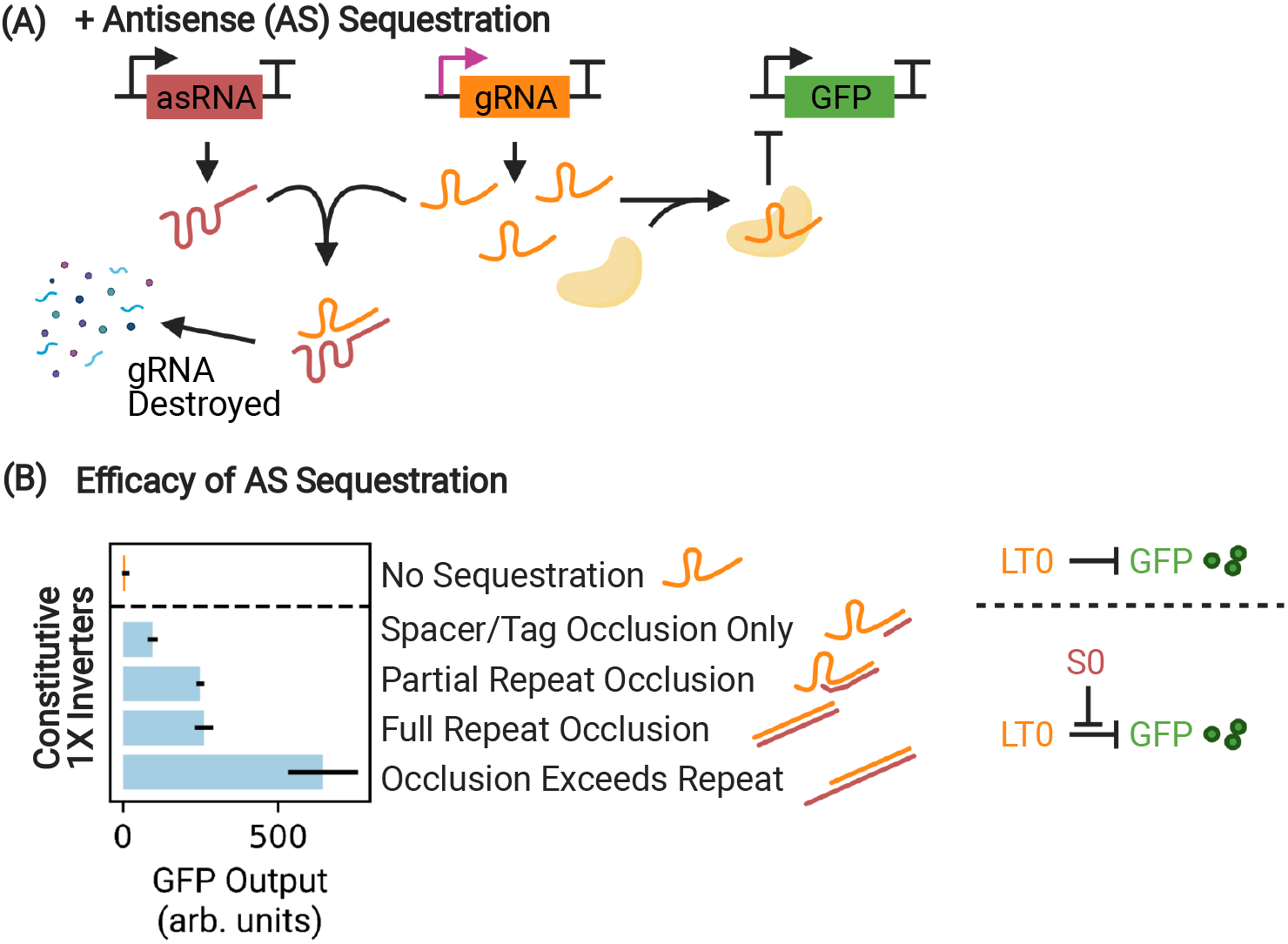
Efficacy of gRNA sequestration measured via interference with a 1X inverter. (A) By soaking up and destroying leaked gRNA transcripts using a matching asRNA sequence, the upstream circuit leak which limits circuit dynamic range can be nullified. (B) Constitutive expression of a 1X inverter (orange) produces cells which are white, as GFP expression is suppressed by dCas binding. By occluding portions of the gRNA (occluding only the spacer/tag, partial or full occlusion of the repeat, and occlusion which exceeds the repeat sequence, light blue), there is a demonstrable difference in sequestration efficacy as a function of interference with the function of CRISPRi, which increases GFP output. Occlusion of the complete gRNA sequence, exceeding the full length of the repeat, results in the most effective sequestration. Ultimately, partial repeat occlusion is used in all subsequent experiments in order to minimize potential non-orthogonality with asRNAs intended to target different gRNAs. Differences in GFP output are measured during exponential growth. The HFQ recruitment tag on the asRNA is not depicted for clarity.

CRISPRi inverters are extremely sensitive to input leak, as the presence of just a few leaked gRNAs may bind to otherwise unoccupied dCas12a and persistently repress their targets until diluted away. As a result, performance of a 1X inverter plummets with respect to a control (constitutive expression of GFP with controlled compositional context, Fig. 3A), especially during stationary expression. Here, the 1X inverter is limited such that only 45.3% of the expected range is covered (orange, Fig. 3A,B, with respect to the dotted green control). During exponential growth, this effect is less extreme, as a basic 1X inverter (orange, Fig. S3) covers 75.0% of the expected maximal dynamic range when compared to the green control. In both instances, poor performance is driven by low maximal GFP expression in the low induction condition. As expected, the basic inverter shows very low although nonzero output leak with respect to the background (3.3% of the maximal inverter OFF state during exponential growth, 3.7% during stationary expression, Figs. 3B and S3).

**Figure 3:**
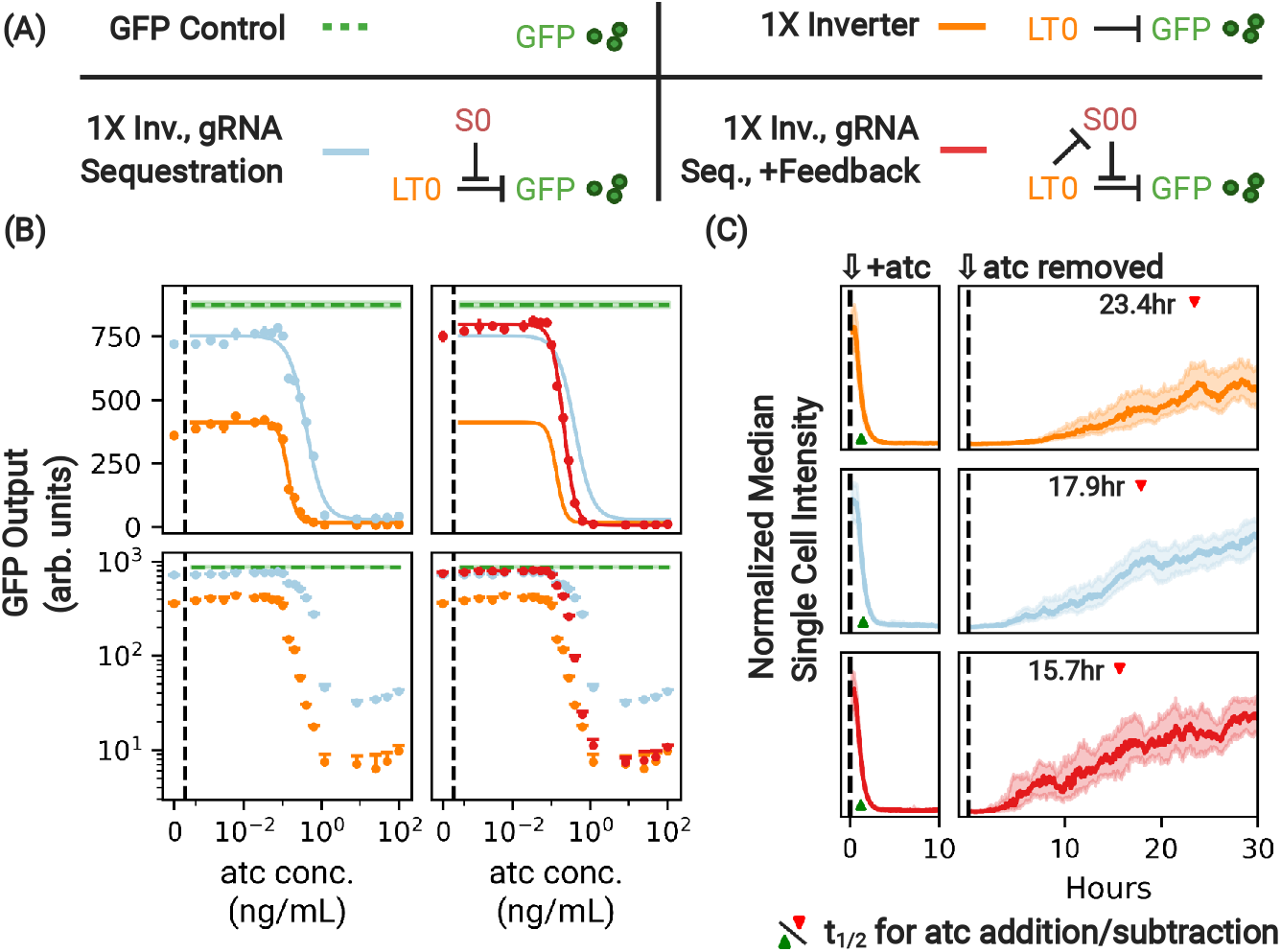
Antisense sequestration of gRNA increases dynamic range of a 1X inverter. (A) A control and variants of the 1X inverter with gRNA sequestration designed to have the same compositional context (see Supplementary Fig. S2). (B) During stationary expression, absolute dynamic range of the basic 1X inverter (orange) is greatly limited by circuit leak, which reduces GFP output with respect to the expected maximum (dotted green) when atc concentration is low. Antisense sequestration of the gRNA via S0 (light blue) acts to suppress CRISPRi repression, expanding the dynamic range of the circuit. However, this comes at the cost of suboptimally higher expression at high induction, evident in logspace. The addition of the feedback mechanism (red) suppresses production of the asRNA when gRNA production is high, maintaining a high dynamic range while nullifying the unwanted impacts of sequestration at high atc concentrations. In linear space displayed error bars are ± 1 standard deviation from threefold biological replicates. Performance is shown relative to the performance of a GFP control with the same compositional context arrangement of nodes (dashed green line) and the basic 1X inverter (orange). For these and all subsequent experiments, dCas12a and tetR are expressed constitutively in the genome. (C) The same constructs, this time under the addition and subtraction of atc in a microfluidic chamber. The presence of antisense sequestration (light blue) speeds circuit response under atc removal (dCas derepression, 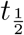 indicated with a red caret) at the cost of some speed in repression (Table 1). Use of the dCas regulatory feedback restores the speed of repression while maintaining improved speed of derepression. Traces are median intensity of single cells across all microfluidic channels. Shaded regions indicate ±1 quartile.

We next sought to determine if antisense sequestration (light blue, Figs. 3A,B) had a substantial effect on circuit performance. Interestingly, we found that antisense sequestration significantly improves the performance with respect to the basic 1X inverter, especially during stationary expression where the dynamic range of the circuit increases from 45.3% to 82.8% of the expected range. During exponential growth, however, the overall dynamic range is relatively unaffected (Fig. S3). Thus, the effect of asRNA during stationary expression is to translate the entire induction curve towards higher levels of GFP production, consistent with the expectation that gRNAs are sequestered at all levels of atc induction.

### Utilization of positive feedback reduces output leak caused by gRNA sequestration

While the absolute dynamic range of a 1X inverter with input leak is vastly improved by the use of antisense sequestration, this comes at some significant cost (first column of Figs. 3B, S3): at high levels of induction, the presence of antisense sequestration also increases output leak, thereby reducing CRISPRi effectiveness. It is important that we mitigate this leak, especially for the instance where the node output is the circuit output (i.e. GFP, rather than another processing node). CRISPRi’s inherently programmable mechanism of action and compact regulatory footprint gives us a way to introduce feedback into the system that can reduce output leak while preserving the benefits of antisense sequestration at low atc induction. Specifically, we implement positive feedback by having the 1X inverter regulate the production of its own antisense sequestration RNA (Fig. 3A, gRNA sequestration + feedback). This system creates a positive feedback mechanism which reinforces sequestration when it is desirable and suppresses it when it is not.

Fig. 3 shows how the use of regulatory feedback entirely removes the output leak penalty introduced by antisense sequestration during stationary expression (red curve). Output leak is reduced to a level comparable to that of the original 1X inverter. Further, the absolute range of the circuit (90.5% of the expected maximal range) is retained, more than doubling the range of the original inverter. Essentially, the regulatory feedback module allows us to use antisense sequestration to control input leak, without affecting the output leak.

However, since the reduction of output leak is less dramatic during exponential growth (Fig. S3), these results support the hypothesis that circuit performance is driven by cellular division time. When cells are rapidly dividing, dCas12a is kicked off of its target by the DNA replication machinery at a higher rate, which in turn impedes complete repression of asRNA in the high atc state. However, when replication rates are low, the system reaches an equilibrium where asRNA production is kept at a low enough level so as to totally restore the full repression of the original inverter. We also attempted to minimize output leak by sequestering mRNA (Figs. S4, S5). This was less successful owing to greatly slowed circuit dynamics and increased noise when used in conjunction with gRNA sequestration and is discussed in the supplementary text.

### Sequestration with regulatory feedback speeds derepression at no cost to repression speed

Although population-wide induction dynamics measured using microplate fluorescence allow us to study equilibrium expression in high density culture, they give us a limited ability to measure alterations of the induction dynamics due to antisense sequestration. Thus, to more precisely understand how asRNA sequestration interacts with our 1X inverter variants, we used a ‘mother machine’ device (Methods, Figs. S8, S9, Movie S1) in order to track the expression level of individual cells in response to induction and repression of the 1X inverter.

We first considered the impact of gRNA sequestration of the repression dynamics when cells are exposed to atc, and on de-repression timescales when atc is removed from the system. In agreement with previous authors, ^25^ we find that the use of antisense sequestration speeds up de-repression, reducing *t*_1/2_ by 24% from 23 to 17 hours (Fig. 3C, noting the position of the red caret, and Movie S1). However, the use of gRNA sequestration increases the response time following the addition of inducer by 15%, suggesting that antisense sequestration may interfere with repression dynamics, even when levels of gRNA are high.

We next investigated whether regulatory feedback further improves the induction/repression dynamics. Fig. 3C shows that gRNA sequestration further reduces the de-repression time when atc is removed, decreasing it by 33% with respect to the original inverter. In addition, the use of the positive feedback mechanism completely restores the induction response time and appears to nullify the increase in response time associated with gRNA sequestration. Thus, our single-cell results shows that a circuit that contains both antisense sequestration and regulatory feedback displays the highest dynamics range and responds up to 33% faster than a “vanilla” 1X inverter.

### Use of antisense sequestration partially restores the logical behavior of a double inverter

To evaluate whether antisense sequestration also improves the dynamic range and response time of multi-layered circuits (i.e. circuits where the input of one logic node is used as the input of another logic node), we constructed a double inverter using the same basic approach to node arrangement as before (alternating logic and sinker nodes Fig. 3A). Other authors have previously observed signal loss in multiple inverters, ^13,18^ and such a result is expected due to dCas leak sensitivity. ^22^ The double inverter, while in principle a simple circuit, is an excellent testbed for measuring how our antisense sequestration system alters circuit performance when nodes are used in series.

We first observe that the performance of the vanilla double inverter is extremely poor during exponential growth, covering only 10.8% of the expected maximal range with respect to the control (Fig. S3, blue) and vastly underperforms the single inverter. Performance of the double inverter does recover during stationary expression, although the maximal expression of a 2X inverter is comparable to that of the single inverter but still underperforms the expected maximal expression (Fig. 4B, in blue).

**Figure 4:**
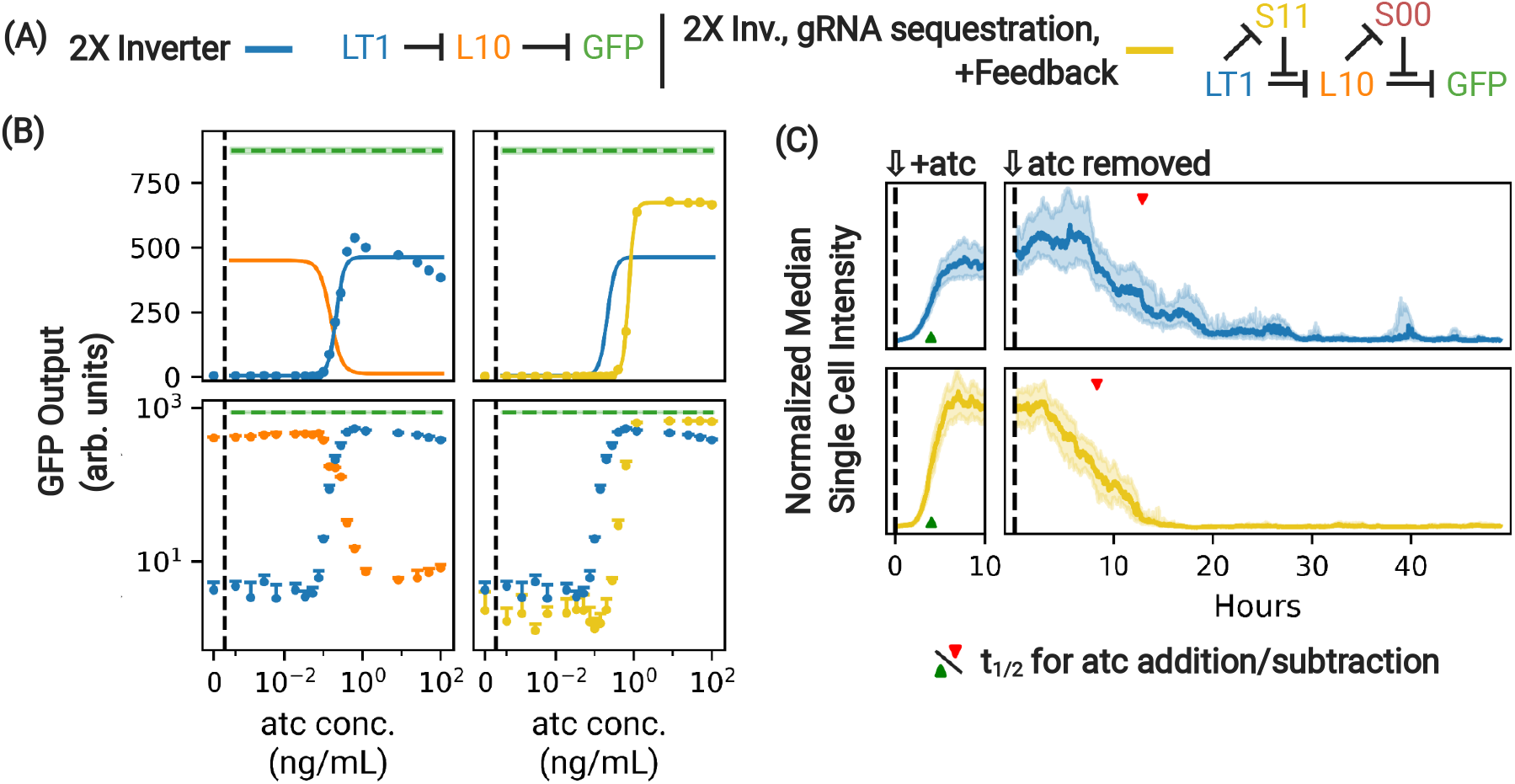
Antisense sequestration of gRNA partially restores the dynamic range of a 2X inverter. (A) Two variants of the 2X inverter controlled to have the same compositional context. (B) Antisense sequestration of gRNA of the 2X inverter (yellow) partially restores the dynamic range by expanding the range of expression in both ON and OFF states, compared to the basic 2X inverter with no sequestration (blue). In linear space displayed error bars are ±1 standard deviation from threefold biological replicates. Performance is compared to the same GFP control (dotted green) used in Fig. 3. Due to the extremely slow equilibration time of the 2X inverter these constructs have been run for additional time in order to be allowed to reach equilibrium (see Methods). (C) The same constructs, this time under the addition and subtraction of atc in a microfluidic chamber. The presence of antisense sequestration speeds circuit response under atc addition and removal with respect to the basic 2X inverter. Traces are median intensity of single cells across all microfluidic channels. Shaded regions indicate ±quartile.

Poor performance in a double inverter is expected to be the result of output leak in the first node without inducer, which in turns drives leaky expression of the second node when it should normally be turned off. As such, we next sought to determine whether the addition of antisense sequestration with feedback, our best-performing system for the 1X inverter, could improve a basic 2X inverter (Fig. 4). Fig. 4B shows that antisense sequestration restores a significant fraction of the dynamic range by expanding the span of expression at both low and high levels of induction. While the absolute dynamic range during exponential growth (42.1%) remains low, its dynamic range is nearly quadrupled compared to the vanilla 2X inverter (Fig. S7).

In addition, single-cell measurements in Fig. 4C shows that the use of sequestration significantly speeds the response of the 2X inverter circuit under atc removal, reducing *t*_1/2_ by 36%, without significantly affecting its performance under atc addition (Table 1). Therefore, despite the double inverter being extremely susceptible to slow dCas12a dynamics due to a large population of programmed dCas12a needs to be replaced in order to reach equilibrium, our single-cell results show that antisense sequestration and regulatory feedback significantly improve both the dynamic range of and response time of 2X inverter when adding or removing the inducer.

**Table 1:** Circuit response to induction as observed via microfluidics. Circuit response time *t*_1/2_ is observed in response to atc addition and removal. *t*_1/2_ is calculated using a spline fit to the median induction curve (see Figs. S10, S11).

| Circuit | $t_{\frac{1}{2}}$ (response to atc addition, min) | $t_{\frac{1}{2}}$ (response to atc removal, min) |
| --- | --- | --- |
| 1X Inverter | 76.8 | 1405.1 |
| 1X Inverter + gRNA Sequestration | 89.0 | 1072.4 |
| 1X Inverter + gRNA Sequestration, Feedback | 72.7 | 940.4 |
| 1X Inverter + mRNA Sequestration, Feedback | 68.8 | 2902.5 |
| 1X Inverter + gRNA/mRNA Sequestration, Feedback | 68.2 | 1219.5 |
| 2X Inverter | 239.5 | 772.3 |
| 2X Inverter + gRNA Sequestration, Feedback | 242.1 | 498.1 |

## Discussion

The use of dCas proteins as programmable repressors holds great promise in synthetic biology, given that they are effective, highly engineerable, and orthogonal. However, these nucleases did not evolve for the purpose of transcriptional repression and suffer as an all-purpose transcription factor, particularly due to sensitivity to leak. Inheritance of leak between upstream and downstream nodes drives poor performance in dCas-based systems, causing oscillators to not oscillate and toggle switches to not toggle.^22^ Hence, a CRISPRi-based system which improves leak tolerance could fix these problems without sacrificing programmability or orthogonality. Further, increasing the effectiveness of dCas-based transcriptional regulators allows us to free up the use of a diverse but limited set of inducible sensors^4^ for sensing rather than internal logic processing components.

Our system improves on suboptimal performance of CRISPRi-based circuits by dealing with circuit leak directly by removing leaked transcripts from the system, similar to ‘sponge sites’ present in natural systems which sequester excess transcription factors via DNA sites in the genome^28–31^ or RNAs.^32^ Because dCas12a associates functionally irreversibly^33^ with its target until displaced by DNA replication, just a few leaked gRNA transcripts can cause significant repression. It is advantageous to regulate gRNA directly, as RNA-based sequestration benefits from a separation of timescales, since gRNA and asRNA diffusion is a fast process that should equilibrate before the demonstrably slow dCas search mechanic.^34^ Further, in the context of CRISPRi-based gene circuits, dCas is physiologically expensive and its maximum concentration is a limiting factor^18^ and nucleases such as dCas9 are even toxic in some bacteria when highly expressed.^35^ By contrast, RNA transcripts are physiologically inexpensive to produce and destroy, unlike dCas proteins, a costly resource.

Dual CRISPRi/antisense RNA (asRNA) elements have been created previously as a means to more rapidly ‘de-repress’ CRISPRi nodes,^25,36^ counteracting excessive memory of initial dCas repression. Microfluidic study allows us to study the performance of cells over long periods of time under constant growth conditions and variable exposure to induction. Our results show that dCas-based regulatory feedback can counteract the slowing of dCas repression by antisense sequestration without cost to beneficial speeding of derepression. Ultimately, we show that a 2X inverter that uses asRNA to limit the impact of transcription leaks responds far more quickly to atc removal.

Remaining challenges include the sensitivity to growth phase and the intrinsic complexity of adding an additional layer of feedback to CRISPRi. Recent work on this subject has often only considered expression during exponential growth stages. However, we believe that it is important to consider the performance of dCas circuits in dynamic growth conditions and at high densities, given the importance of host/circuit interactions in the performance of synthetic gene circuits. ^37,38^ During stationary-phase growth with constant protein production,^39^ more dCas proteins find their cognate target, strengthening repression and thus leak sensitivity. Ultimately, it is important to understand how these circuits perform and can be improved under conditions that are more relevant in industrial application.^40^

Our solution improves the performance of dCas circuits without sacrificing the attribute which makes them so desirable: programmability. This does, however, come at the cost of increased complexity. Despite this, we believe that merging antisense sequestration with CRISPRi is necessary in order to mitigate potentially crippling issues with CRISPRi and other dCas-based transcription factors. Further, the use of feedback in essential for robustness in engineered circuits. ^41^ Especially with further advancement in the cell-free assembly of large plasmids (e.g. OriCiro^42^), it will only become easier to create larger systems of dCas synthetic gene networks.

In this work, we have demonstrated that antisense RNA-based sponge sites can be used to reduce leak sensitivity for CRISPRi-based gene circuits, particularly during slow growth when leaked gRNA transcripts drive unintended repression and reduced dynamic range. While this work only explores the impacts of sequestration on NOT gate CRISPRi repressors, in principle this technique could also be used with CRISPRa.^11,12^ Additionally, this work could be combined with dCas degradation, which should in principle further reduce leak sensitivity, ^22^ or dCas self-regulation, which could further fortify circuit performance. ^21^ Overall, our system reduces leak sensitivity in CRISPRi systems, which will help to tap their potential to create complex and engineerable genetic systems that can rival those found in nature.

## Materials and Methods

### Plasmid assembly

The original CRISPRi and sinker nodes were ordered as gene blocks from IDT and inserted into pUC19 for ease of modification. Site-directed mutagenesis to modify the sequences was done exclusively using NEB’s Q5 Hot Start and NEB’s KLD (kinase, ligase, and dpn1 digestion) prior to transformation. Verification of correct sequences was done using Sanger sequencing via Cornell’s Genomics Facility. Modification of individual nodes was completed in pUC19 plasmid before digestion and insertion into the main experimental plasmid which contains the circuit. Assembly of multinode circuits was done using standard molecular biology techniques for *E. coli* using restriction enzyme digestion with AarI (Thermo Fisher) and BsaI-HFv2 (NEB). Ligation of digested components was done using NEB’s Instant Ligase. Colony PCR using NEB’s Taq polymerase was used to check for successful node insertion.

All cloning was done in NEB Stable in order to minimize possible recombination events due to the use of repetitive sequences. Cells used for cloning were cultured using liquid TB (Terrific Broth, VWR) prepared using the manufacturer’s instructions. Plasmids were maintained using antibiotics as appropriate (Kanamycin, Chloramphenicol, and Ampicillin). LB (lysogeny broth, VWR) agar plates were used as a solid medium.

FnCas12a is made catalytically dead via D917A and E1006A mutations. All GFP sequences have an orthogonal ssrA tag^43^ for degradation, although this was not induced in the course of these experiments. Annotated plasmid sequences used in this study can be found in the Supplementary Materials and are hosted by Benchling.

### Node design

Generally, CRISPRi and antisense sequestration nodes are designed to be a standard part, with comparable expression strength. The design of nodes used in this system is illustrated in Supplementary Fig. S1. Every gRNA-producing node (including the pTet logic node), termed a ‘logic’ node, and every asRNA-producing node, termed a ‘sinker’ node, shares the same strong promoter with conserved −35 and −10 promoter sites TTGACA and TAAAAT. The output promoter which drives GFP is designed to be weaker (−35 and −10 sites TTGTCA and TAAAAT), as expression of GFP by the strong promoter causes a fitness penalty. All nodes (except those driven by pTet) have a PAM site TTTG necessary for dCas binding for logical control or feedback.

All nodes are inserted in a tandem orientation as indicated in Fig. S1. Annotated versions of the logic node (https://benchling.com/s/seq-JItVzOqT6qBnrCk0aLcn), pTet-driven logic node (https://benchling.com/s/seq-gdhKvLmV6Xs6u2lbNLFb), and sinker node (https://benchling.com/s/seq-s6qBkmmvJ38VzyDX1RHB) are hosted via Benchling. Forty bp randomized tags with fixed GC content are used to produce cognate asRNAs which orthogonally sequester gRNA sequences. Two asRNA sequences exhibited toxic effects when expressed and were not used further in the study (see Supplementary Materials, Sinker Node Archetype). The HFQ tag used to facilitate sRNA sequestration is the micF M7.4 tag from 26.

Throughout these experiments, we have taken care to preserve the compositional context of circuits that we seek to compare. We anecdotally observed that the order of nodes can significantly impact the performance of these gene circuits. None of these variants significantly outperform the original node ordering.

### Microplate fluorescence assays

All measures of fluorescence are conducted using the GL002 strain unless otherwise noted. GL002 is a variant of the F3 strain with genomically integrated ^44^ expression of tetR, lacI, and dCas12a. The F3 strain (Wakamoto Lab, University of Tokyo) contains knockouts of fliC, fimA, and flu which decrease cell aggregation and adhesion to surfaces.^45^ Cells were made electrocompetent, electroporated at 1800V (BTX ECM399), and recovered in NEB stable media prior to plating for colony selection. Measures of fluorescence where tetR and dCas12a are expressed in a second plasmid (Fig. 2) are done using the F3 strain.

The experimental procedure was as follows. Electrocompetent GL002 cells are electroporated with the pSC101 plasmid containing a complete circuit and plated as described above. After colonies are visible on the next day, three different colonies are selected to inoculate three different 10mL cell culture tubes, each with 2mL H media with antibiotic (Kanamycin) and grown to saturation overnight for 18-19 hours in a shaker held at 37 °C. 100μL of this culture is then used to inoculate a tray containing 2mL H media with antibiotic which is shaken until homogenous. 1 μL volumes are taken from this tray using a multipipettor and used to inoculate wells of a 96-well plate (VWR, catalog number 10062-900) with 200*μ*L volumes of H media, appropriate antibiotics, and inducer. A sacrificial border of 36 200 μL volumes surrounds the 60 wells used for each experiment on the plate to minimize evaporative losses. Quantitative measurements of fluorescence were made using a Synergy H1 Hybrid Multi-Mode Microplate Reader, produced by BioTek, with temperature held at 37 °C and linear shaking at 10 second intervals. Top and bottom fluorescence measurements and 600 nm absorbance measurements were taken at 3 minute intervals, although only the top measurements are reported here.

In experiments studying the 2X inverter, only 1 μL of overnight culture is used to inoculate the plate, owing to the anticipated extremely long equilibration time under atc addition. All other experimental parameters are held constant.

H media^46^ was used throughout these experiments both because it is sufficiently rich yet minimally autofluorescent, easing microfluidic study. H media is LB without yeast extract - 10 g/L tryptone (BD), 8 g/L NaCl (VWR).

In order to account for small variations in inoculation volume, fluorescence curves are aligned using the absorbance measure. Curves are aligned to the time when they cross an absorbance (corrected for media absorbance) of 0.04, which corresponds to a standard optical density OD600 of 0.16. Background fluorescence was subtracted, calculated via the fluorescence of GL002 cells which contained plasmid w37, which contains the pSC101 origin but lacks GFP. Fluorescence reads are smoothed using SciPy’s implementation of the Wiener filter.

Atc (anhydrotetracycline, Alfa Aesar), used for pTet induction, was kept at a stock concentration of 100 ng/1 μL in a 50% ethanol solution and protected from light.

### Induction analysis

Induction curves are taken at slices in time with respect to the time when the alignment OD is reached (see Microplate fluorescence assays). The times are 60 and 500 minutes with respect to the alignment time for exponential and stationary expression, respectively.

Induction curves are fit to a Hill function of form:

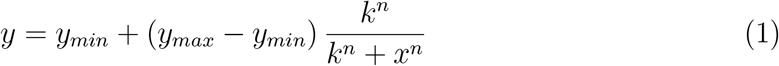

The values of *y*_*min*_ and *y*_*max*_ are used for calculations of absolute dynamic range and fold change.

### Microfluidic experimental design

Dynamic induction experiments were performed in a microfluidic device with chambers of two sizes, with lateral dimensions of L x W = 40.5 × 7.1 μm^2^ and L x W = 35 × 7.1 μm^2^. Only data from the shorter chambers are reported in this study for clarity.

As with prior experiments, plasmids containing the circuit of interest were electroporated into the GL002 strain of *E. coli* and grown on a plate overnight. The following morning cells are inoculated and grown in 3mL of H media with kanamycin to an OD of 0.5, concentrated by centrifugation, and pipetted into the plasma cleaned microfluidic device. The device was then placed in an imaging set up with temperature held at 37 °C. A bottle of fresh media (H media + 1x kanamycin + bovine serum albumin (100mg/L) ± atc) pressurized to 5 psi is used to deliver a constant flow of fresh media (100 mL/day) to the cells.

Cells are introduced and allowed to populate microfluidic chambers. For all constructs, cells are inoculated in the absence of inducer and allowed to equilibrate. After the inducer is added, the system is allowed to equilibrate again, and then the inducer is taken away so that we can measure the return to the initial state (Fig. S9). The full induction is run over the course of at least 80 hours in order to allow the system to reach equilibrium initially, post atc induction, and allow us to measure the response time when atc is removed. Response time *t*_1/2_ is defined as the time taken to reach halfway between the equilibrium minimum and maximum levels of expression, in linear space. Expression levels (as observed in the microfluidic device) and the measured response time are included in Table 1.

Single cells are resolved by epi-fluorescence microscopy of sf-GFP with a 100x, 1.4 NA apochromat Leica objective. We typically observe the circuits in its non-induced state overnight, then add atc to induce the circuit for 23 hours ±15 min and finally record the recovery after removal of atc for 40 to 60 hours. We monitor 20 to 40 chambers in parallel in a given experiment.

## Notes

### Competing Interest Statement

The authors have declared no competing interest.

## References

(1) Potvin-Trottier, L.; Lord, N. D.; Vinnicombe, G.; Paulsson, J. Synchronous longterm oscillations in a synthetic gene circuit. Nature 2016, 538, 514–517.

(2) Gardner, T. S.; Cantor, C. R.; Collins, J. J. Construction of a genetic toggle switch in Escherichia coli. Nature 2000, 403, 339–342.

(3) Nielsen, A. A. K.; Voigt, C. A. Multi-input CRISPR/Cas genetic circuits that interface host regulatory networks. Molecular Systems Biology 2014, 10, 763.

(4) Meyer, A. J.; Segall-Shapiro, T. H.; Glassey, E.; Zhang, J.; Voigt, C. A. Escherichia coli “Marionette” strains with 12 highly optimized small-molecule sensors. Nature Chemical Biology 2019, 15, 196–204.

(5) Chen, Y.-J.; Liu, P.; Nielsen, A. A. K.; Brophy, J. A. N.; Clancy, K.; Peterson, T.; Voigt, C. A. Characterization of 582 natural and synthetic terminators and quantification of their design constraints. Nature Methods 2013, 10, 659–664.

(6) Qi, L. S.; Larson, M. H.; Gilbert, L. A.; Doudna, J. A.; Weissman, J. S.; Arkin, A. P.; Lim, W. A. Repurposing CRISPR as an RNA-Guided Platform for Sequence-Specific Control of Gene Expression. Cell 2013, 152, 1173–1183.

(7) Larson, M. H.; Gilbert, L. A.; Wang, X.; Lim, W. A.; Weissman, J. S.; Qi, L. S. CRISPR interference (CRISPRi) for sequence-specific control of gene expression. Nature Protocols 2013, 8, 2180–2196.

(8) Kim, S. K.; Kim, H.; Ahn, W.-C.; Park, K.-H.; Woo, E.-J.; Lee, D.-H.; Lee, S.-G. Efficient Transcriptional Gene Repression by Type V-A CRISPR-Cpf1 from Eubacterium eligens. ACS synthetic biology 2017, 6, 1273–1282.

(9) Specht, D. A.; Xu, Y.; Lambert, G. Massively parallel CRISPRi assays reveal concealed thermodynamic determinants of dCas12a binding. Proceedings of the National Academy of Sciences 2020, 117, 11274–11282.

(10) Jusiak, B.; Cleto, S.; Perez-Piñera, P.; Lu, T. K. Engineering Synthetic Gene Circuits in Living Cells with CRISPR Technology. Trends in Biotechnology 2016, 34, 535–547.

(11) Liu, Y.; Wan, X.; Wang, B. Engineered CRISPRa enables programmable eukaryote-like gene activation in bacteria. Nature Communications 2019, 10, 3693.

(12) Fontana, J.; Dong, C.; Kiattisewee, C.; Chavali, V. P.; Tickman, B. I.; Carothers, J. M.; Zalatan, J. G. Effective CRISPRa-mediated control of gene expression in bacteria must overcome strict target site requirements. Nature Communications 2020, 11, 1618.

(13) Didovyk, A.; Borek, B.; Hasty, J.; Tsimring, L. Orthogonal Modular Gene Repression in Escherichia coli Using Engineered CRISPR/Cas9. ACS synthetic biology 2016, 5, 81–88.

(14) Cress, B. F.; Jones, J. A.; Kim, D. C.; Leitz, Q. D.; Englaender, J. A.; Collins, S. M.; Linhardt, R. J.; Koffas, M. Rapid generation of CRISPR/dCas9-regulated, orthogonally repressible hybrid T7-lac promoters for modular, tuneable control of metabolic pathway fluxes in Escherichia coli. Nucleic Acids Research 2016, 44, 4472–4485.

(15) Santos-Moreno, J.; Tasiudi, E.; Stelling, J.; Schaerli, Y. Multistable and dynamic CRISPRi-based synthetic circuits. Nature Communications 2020, 11, 2746.

(16) Kuo, J.; Yuan, R.; Sánchez, C.; Paulsson, J.; Silver, P. A. Toward a translationally independent RNA-based synthetic oscillator using deactivated CRISPR-Cas. Nucleic Acids Research 2020, 48, 8165–8177.

(17) Henningsen, J.; Schwarz-Schilling, M.; Leibl, A.; Gutierrez, J.; Sagredo, S.; Simmel, F. C. Single Cell Characterization of a Synthetic Bacterial Clock with a Hybrid Feedback Loop Containing dCas9-sgRNA. ACS Synthetic Biology 2020, 9, 3377–3387.

(18) Zhang, S.; Voigt, C. A. Engineered dCas9 with reduced toxicity in bacteria: implications for genetic circuit design. Nucleic Acids Research 2018, 46, 11115–11125.

(19) Jayanthi, S.; Nilgiriwala, K. S.; Del Vecchio, D. Retroactivity Controls the Temporal Dynamics of Gene Transcription. ACS Synthetic Biology 2013, 2, 431–441.

(20) Brophy, J. A. N.; Voigt, C. A. Principles of genetic circuit design. Nature Methods 2014, 11, 508–520.

(21) Huang, H.-H.; Bellato, M.; Qian, Y.; Cárdenas, P.; Pasotti, L.; Magni, P.; Del Vecchio, D. dCas9 regulator to neutralize competition in CRISPRi circuits. Nature Communications 2021, 12, 1692.

(22) Clamons, S. E.; Murray, R. M. Modeling Dynamic Transcriptional Circuits with CRISPRi. bioRxiv 2017, 225318.

(23) Zetsche, B.; Gootenberg, J. S.; Abudayyeh, O. O.; Slaymaker, I. M.; Makarova, K. S.; Essletzbichler, P.; Volz, S. E.; Joung, J.; van der Oost, J.; Regev, A.; Koonin, E. V.; Zhang, F. Cpf1 Is a Single RNA-Guided Endonuclease of a Class 2 CRISPR-Cas System. Cell 2015, 163, 759–771.

(24) Lutz, R.; Bujard, H. Independent and tight regulation of transcriptional units in Escherichia coli via the LacR/O, the TetR/O and AraC/I1-I2 regulatory elements. Nucleic Acids Research 1997, 25, 1203–1210.

(25) Lee, Y. J.; Hoynes-O’Connor, A.; Leong, M. C.; Moon, T. S. Programmable control of bacterial gene expression with the combined CRISPR and antisense RNA system. Nucleic Acids Research 2016, 44, 2462–2473.

(26) Hoynes-O’Connor, A.; Moon, T. S. Development of Design Rules for Reliable Antisense RNA Behavior in E. coli. ACS Synthetic Biology 2016, 5, 1441–1454.

(27) Møller, T.; Franch, T.; Højrup, P.; Keene, D. R.; Bächinger, H. P.; Brennan, R. G.; Valentin-Hansen, P. Hfq: A Bacterial Sm-like Protein that Mediates RNA-RNA Interaction. Molecular Cell 2002, 9, 23–30.

(28) Lee, T.-H.; Maheshri, N. A regulatory role for repeated decoy transcription factor binding sites in target gene expression. Molecular Systems Biology 2012, 8, 576.

(29) Brewster, R. C.; Weinert, F. M.; Garcia, H. G.; Song, D.; Rydenfelt, M.; Phillips, R. The Transcription Factor Titration Effect Dictates Level of Gene Expression. Cell 2014, 156, 1312–1323.

(30) Kemme, C. A.; Esadze, A.; Iwahara, J. Influence of Quasi-Specific Sites on Kinetics of Target DNA Search by a Sequence-Specific DNA-Binding Protein. Biochemistry 2015, 54, 6684–6691.

(31) Kemme, C. A.; Nguyen, D.; Chattopadhyay, A.; Iwahara, J. Regulation of transcription factors via natural decoys in genomic DNA. Transcription 2016, 7, 115–120.

(32) Morriss, G. R.; Cooper, T. A. Protein sequestration as a normal function of long noncoding RNAs and a pathogenic mechanism of RNAs containing nucleotide repeat expansions. Human genetics 2017, 136, 1247–1263.

(33) Strohkendl, I.; Saifuddin, F. A.; Rybarski, J. R.; Finkelstein, I. J.; Russell, R. Kinetic Basis for DNA Target Specificity of CRISPR-Cas12a. Molecular Cell 2018, 71, 816–824.e3.

(34) Jones, D. L.; Leroy, P.; Unoson, C.; Fange, D.; Ćurić, V.; Lawson, M. J.; Elf, J. Kinetics of dCas9 target search in Escherichia coli. Science 2017, 357, 1420–1424.

(35) Cho, S.; Choe, D.; Lee, E.; Kim, S. C.; Palsson, B.; Cho, B.-K. High-Level dCas9 Expression Induces Abnormal Cell Morphology in Escherichia coli. ACS synthetic biology 2018, 7, 1085–1094.

(36) Mückl, A.; Schwarz-Schilling, M.; Fischer, K.; Simmel, F. C. Filamentation and restoration of normal growth in Escherichia coli using a combined CRISPRi sgRNA/antisense RNA approach. PLOS ONE 2018, 13, e0198058.

(37) Nikolados, E.-M.; Weiße, A. Y.; Ceroni, F.; Oyarzún, D. A. Growth Defects and Loss-of-Function in Synthetic Gene Circuits. ACS Synthetic Biology 2019, 8, 1231–1240.

(38) Tan, C.; Marguet, P.; You, L. Emergent bistability by a growth-modulating positive feedback circuit. Nature Chemical Biology 2009, 5, 842–848.

(39) Gefen, O.; Fridman, O.; Ronin, I.; Balaban, N. Q. Direct observation of single stationary-phase bacteria reveals a surprisingly long period of constant protein production activity. Proceedings of the National Academy of Sciences 2014, 111, 556–561.

(40) Moser, F.; Broers, N. J.; Hartmans, S.; Tamsir, A.; Kerkman, R.; Roubos, J. A.; Bovenberg, R.; Voigt, C. A. Genetic Circuit Performance under Conditions Relevant for Industrial Bioreactors. ACS Synthetic Biology 2012, 1, 555–564.

(41) Segall-Shapiro, T. H.; Sontag, E. D.; Voigt, C. A. Engineered promoters enable constant gene expression at any copy number in bacteria. Nature Biotechnology 2018, 36, 352–358, Number: 4 Publisher: Nature Publishing Group.

(42) Hasebe, T.; Narita, K.; Hidaka, S.; Su’etsugu, M. Efficient Arrangement of the Replication Fork Trap for In Vitro Propagation of Monomeric Circular DNA in the Chromosome-Replication Cycle Reaction. Life (Basel, Switzerland) 2018, 8.

(43) Cameron, D. E.; Collins, J. J. Tunable protein degradation in bacteria. Nature biotechnology 2014, 32, 1276–1281.

(44) St-Pierre, F.; Cui, L.; Priest, D. G.; Endy, D.; Dodd, I. B.; Shearwin, K. E. One-Step Cloning and Chromosomal Integration of DNA. ACS Synthetic Biology 2013, 2, 537–541.

(45) Lambert, G.; Kussell, E. Memory and Fitness Optimization of Bacteria under Fluctuating Environments. PLOS Genetics 2014, 10, e1004556.

(46) Elbing, K.; Brent, R. Recipes and tools for culture of Escherichia coli. Current protocols in molecular biology 2019, 125, e83.

