## Supplementary Materials for "Overcoming leak sensitivity in CRISPRi circuits using antisense RNA sequestration and regulatory feedback"

March 24, 2022

#### 1 Supplementary Text

##### 1.1 Combined mRNA and gRNA sequestration maximizes fold change and speeds dynamics of the 1X inverter at the cost of single-cell noise

While output leak is almost completely suppressed by the feedback mechanism at long times during stationary expression, there is still significant leak during exponential growth as a result of antisense sequestration (Fig. S3). We tried to see if we could reduce this output leak even further by sequestering the mRNA output. Design of this asRNA is simple and is made to completely occlude the ribosomal binding site (RBS) for GFP translation (Fig. S1C,G). The regulatory feedback mechanism described in the main text is implemented for the mRNA sequestration using a gRNA sequence expressed on the same transcript as GFP (Fig. S1A), made possible by the fact that Cas12a processes its own gRNA by cleaving upstream of the repeat hairpin. It is important to note, however, that both GFP and the gRNA which suppresses mRNA sequestration are driven by a weaker consensus sequence, and thus the strength of the feedback mechanism is not the same as for the gRNA feedback.

Puzzlingly, at low levels of induction and during stationary expression, expression of GFP exceeds that of the 1X inverter with no mRNA sequestration at low levels of induction (pink curve, Fig. S4). This is an unintuitive result, as one would not initially expect that any amount of mRNA sequestration, even with feedback to control it in the low atc state, would increase overall fluorescence in any condition. However, this is the result of competition between the two gRNAs present in this system. The presence of the second, competing gRNA (light purple, on the same transcript as GFP) competes with would-be leaked transcripts (orange), driving down the efficacy of leaky repression and hence increasing fluorescence. Essentially, we see some 'beneficial' aspect of retroactivity akin to observations by other authors in Ref. 21.

As such, the overall dynamic range of the 1X inverter is increased during stationary expression as the dCas competition effect dominates, increasing absolute dynamic range such that 65.5% of the expected maximum is covered. However, during exponential growth and in line with expectations, mRNA sequestration is the dominant effect, with dynamic range greatly reduced with respect to the original inverter. Slow feedback dynamics mean that retroactivity and leakiness effects dominate during slow post-exponential growth, while mRNA sequestration dominates during faster growth when dCas repression is mitigated by dilution and DNA replication.

As expected, absolute output leak with mRNA sequestration is extremely low, slightly lower than that of the original 1X inverter during all stages of growth. However, the objective of mRNA sequestration is not strictly to reduce the output leakiness of the basic 1X inverter, which was already extremely low. We sought to combine input and output sequestration into a combined system which could balance the benefits of decreased input and output leak (dark purple, Fig. S4A,B). During stationary expression, performance improves such that it has improved dynamic range over the original inverter (76.5% of the expected maximum) and the lowest leak of any inverter that we tested. However, it has reduced dynamic range over the system with gRNA sequestration and feedback alone (red, Fig. 3B). If instead we consider circuit fold change (defining this as $\log_2$ of the Hill function maximum over minimum expression), the 1X inverter with combined mRNA/gRNA sequestration has the highest fold change of any inverter we tested, although only marginally more than the 1X inverter with gRNA sequestration and feedback. During exponential growth, the effect of this combined mRNA and gRNA sequestration unfortunately is to increase both input and output leak with respect to the basic inverter (Fig. S5), resulting in low dynamic range (33.6% of the maximum expected).

As before, we tested these circuits in a microfluidic device in order to study the response dynamics under atc addition and removal (Fig. S4, Table 1). As could be expected, mRNA sequestration has essentially the opposite effect of gRNA sequestration and dramatically slows derepression, increasing $t_{1/2}$ by 106%. Correspondingly, repression is sped slightly, with $t_{1/2}$ decreasing by 10%. Combined mRNA and gRNA sequestration speeds response to changes in induction in both directions, decreasing $t_{1/2}$ by 13% and 11% when atc is removed or added, respectively. However, the competing mechanisms of mRNA and gRNA sequestration considerably increase single-cell variability, producing extremely noisy derepression (Fig. S4C, dark purple).

##### 1.2 Nodes Notation

We use sets of 3-character codes in order to specify nodes and their ordering. This is helpful to notate circuit design and node order as multiple levels of feedback are added. The first character indicates whether the node produces a gRNA (L- logic node) or an asRNA (S - sinker node). The second character indicates the identity of the promoter driving the node, and the third character indicates the identity of its output. Promoters, gRNA, and asRNA pairings are indicated with a number, except T which refers to the inducible pTet promoter. For example, node L10 is driven by promoter 1 and produces a gRNA which targets promoter 0. Node S00 is driven by promoter 0 and produces an asRNA which targets gRNA sequence 0. 'Dud' nodes are nonfunctional stand-ins which express a gRNA or asRNA sequence, as appropriate, but do not target anything in the system. This notation with respect to circuit design is illustrated further in Fig. S1.

#### 2 Supplementary Figures

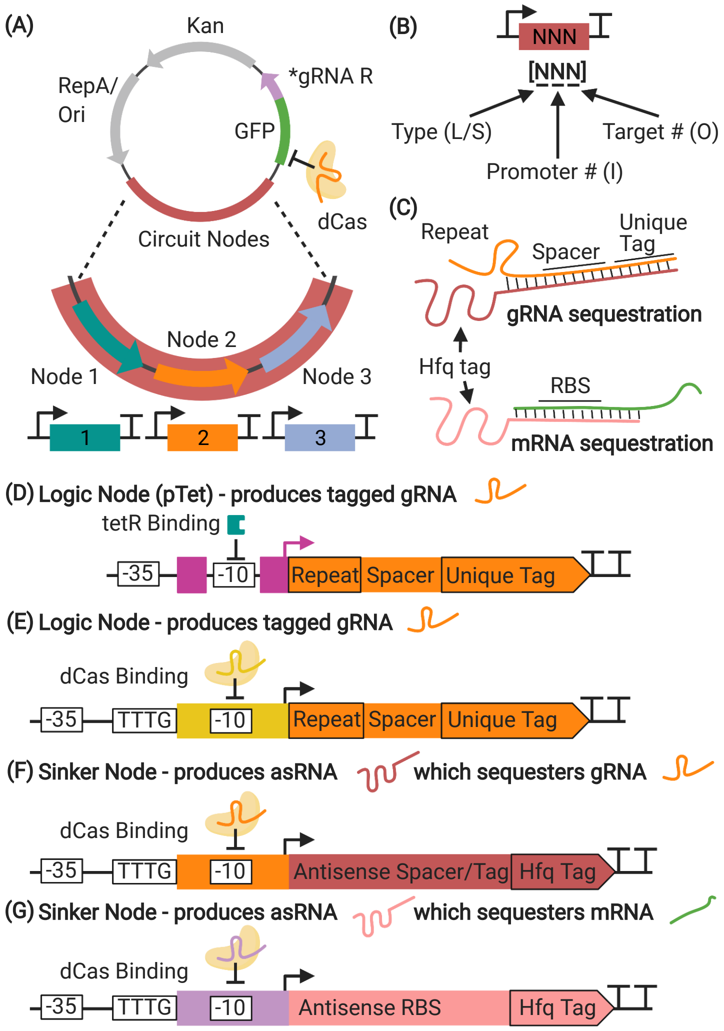


Supplementary Figure S1: **Design of a multinode dual CRISPRi/asRNA system.** (A) Nodes are inserted into a plasmid in tandem with the GFP output. A gRNA which represses mRNA sequestration (asterisked) is present only in constructs with mRNA sequestration. (B) Nodes are notated with a 3-character system designating the node type (logic or sinker), the promoter number, and the output number (either CRISPRi target or asRNA tag). This is useful for specifying node order as circuits get larger and more complex. (C) gRNA sequestration is accomplished using an antisense sequestration sequence which entirely covers the gRNA spacer sequence, an additional 40bp unique tag, and 9bp of the repeat sequence. mRNA sequestration is accomplished by sequestering the mRNA RBS sequence. (D,E,F,G) Each individual node can be repressed for the purposes of logical control or feedback. All nodes have fixed -35 and -10 conserved sequences, although the promoter which drives GFP has a weakened -35 site.

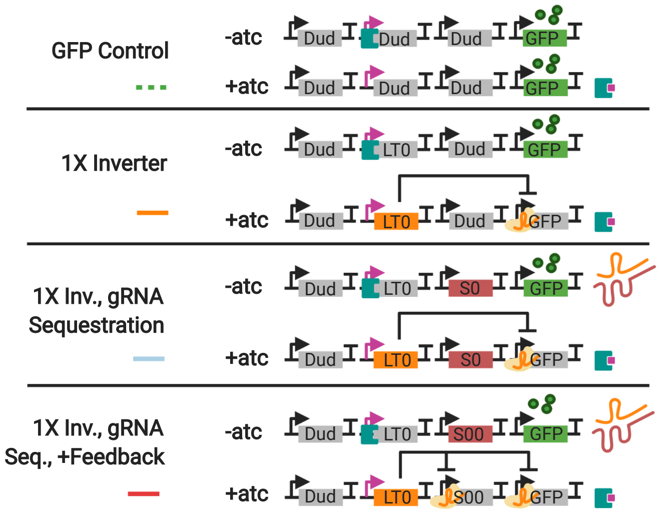


Supplementary Figure S2: **Compositional arrangement of 1X Inverters.** In these experiments the ordering of nodes is fixed. A ‘Dud’ node indicates a nonfunctional stand-in which expresses a gRNA or asRNA sequence, as appropriate, but that does not target anything in the system. The pTet logic node (orange) occurs second, the gRNA sequestration node (red) appears third, followed by the GFP output node. Nodes are notated with a 3-character system designating the node type (logic or sinker), the promoter number, and the output number (either CRISPRi target or asRNA tag), as indicated Supplementary Fig. S1B.

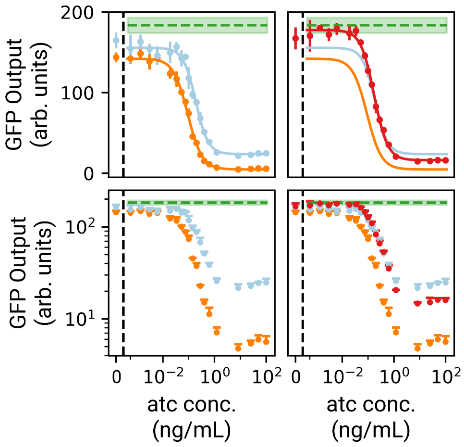


Supplementary Figure S3: **Effect of gRNA sequestration during exponential growth.** Circuit output for 1X inverters without sequestration (orange), with gRNA sequestration (light blue), and with gRNA sequestration and regulatory feedback. Circuits are as diagrammed in Figure 3A. Fit is against a Hill function (see Methods) applied in linear space. In linear space displayed error bars are ±1 standard deviation from threefold biological replicates. Only the upper error bar is displayed in logarithmic space. Performance is shown relative to the performance of a GFP control with the same compositional context arrangement of nodes (dashed green line).

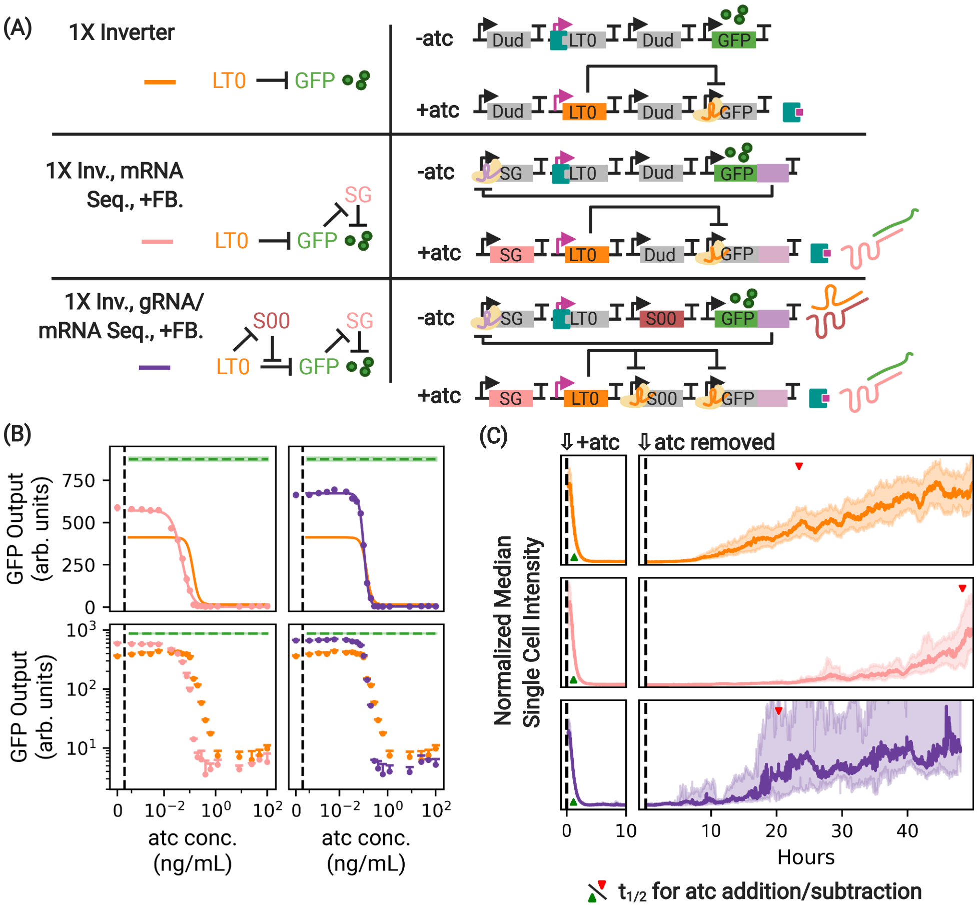

(A)
1X Inverter
LT0
GFP
-atc
+atc
Dud
LT0
GFP
1X Inv., mRNA Seq., +FB.
SG
S00
1X Inv., gRNA/ mRNA Seq., +FB.
(B)
GFP Output (arb. units)
750
500
250
0
atc conc. (ng/mL)
(C)
⇩+atc
⇩atc removed
Normalized Median Single Cell Intensity
Hours
$t_{1/2}$ for atc addition/subtraction

Supplementary Figure S4: **Antisense sequestration of mRNA maximizes dynamic range during stationary expression.** (A) Two additional variants of the single inverter which use mRNA sequestration, compared to a green control and the basic 1X inverter. In these experiments the ordering of nodes is fixed: the mRNA sequestration node (pink) occurs first, the pTet logic node (orange) occurs second, the gRNA sequestration node (red) appears third, followed by the GFP output node. A gray node indicates a nonfunctional stand-in which expresses a gRNA or asRNA sequence, as appropriate, but that does not target anything in the system. In this construct, in order to control expression of the antisense sequence which targets GFP mRNA, a gRNA which targets node SG is expressed on the same transcript as GFP (indicated in light purple). (B) Antisense sequestration of the mRNA via SG acts to suppress excess GFP production. Interestingly, however, the primary effect of mRNA sequestration is not to affect GFP expression at high atc concentration, but rather to increase GFP expression at low atc concentration, an effect of the competition for dCas discussed in the main text. Performance of the 1X inverter with gRNA sequestration (pink) and with gRNA and mRNA sequestration (purple) are shown. Fit is against a Hill function (see Methods) applied in linear space. In linear space displayed error bars are $\pm 1$ standard deviation from threefold biological replicates. Only the upper error bar is displayed in logarithmic space. Performance is shown relative to the performance of a GFP control with the same compositional context arrangement of nodes (dashed green line) and the basic 1X inverter (orange). For these experiments, dCas12a and tetR are expressed constitutively in the genome. (C) The same constructs, this time under the addition and subtraction of atc in a microfluidic chamber. The presence of mRNA sequestration (pink) dramatically slows circuit response under atc removal (dCas derepression) with some advantageous speeding of circuit response under atc addition. Use of combined gRNA and mRNA sequestration restores the speed of derepression while maintaining improved speed of repression. However, this comes at the cost of extreme cell-to-cell variability as expression levels return to the initial state. Traces are median intensity of single cells across all microfluidic channels. Shaded regions indicate $\pm$quartile. Full traces across all microfluidic channels are included in Fig. S9.

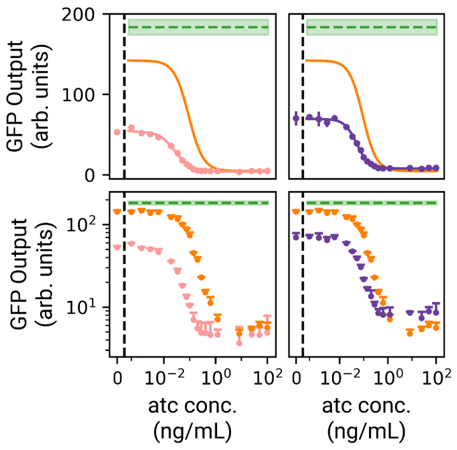


Supplementary Figure S5: **Effect of mRNA sequestration during exponential growth.** Circuit output for 1X inverters without sequestration (orange), with mRNA sequestration and feedback (pink), and with combined mRNA and gRNA sequestration. Circuits are as diagrammed in Fig. S4A. Fit is against a Hill function (see Methods) applied in linear space. In linear space displayed error bars are ± 1 standard deviation from threefold biological replicates. Only the upper error bar is displayed in logarithmic space. Performance is shown relative to the performance of a GFP control with the same compositional context arrangement of nodes (dashed green line).

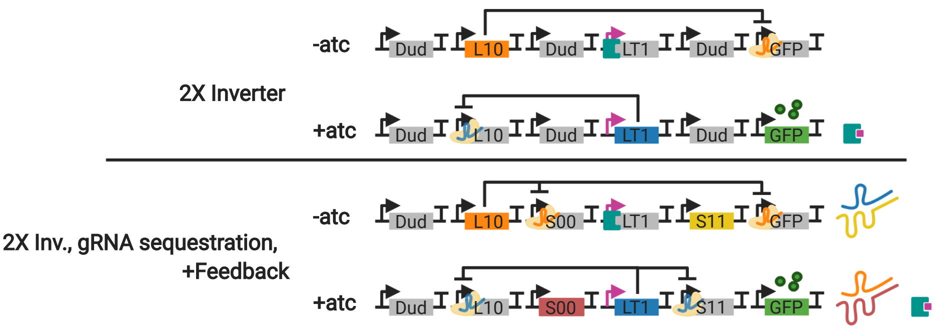


Supplementary Figure S6: **Compositional arrangement of 2X Inverters.** Two variants of the 2X inverter controlled to have the same compositional context. As in the previous figures, Dud nodes indicate a nonfunctional stand-in which expresses a gRNA or asRNA, as appropriate, but that does not target anything in the system.

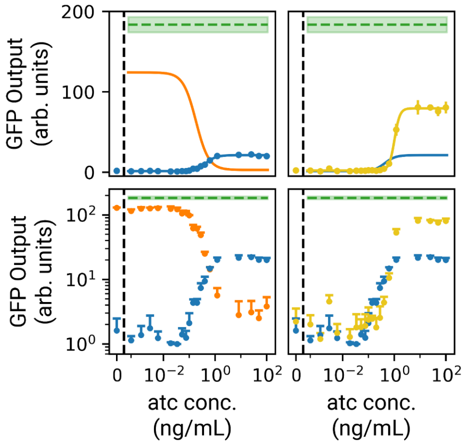


Supplementary Figure S7: **Effect of antisense sequestration on the 2X inverter during exponential growth.** Circuit output for 2X inverters without sequestration (blue) and with gRNA sequestration (yellow), compared against the 1X inverter without sequestration (orange). Circuits are as diagrammed in Fig. 4A. Fit is against a Hill function (see Methods) applied in linear space. In linear space displayed error bars are ±1 standard deviation from threefold biological replicates. Only the upper error bar is displayed in logarithmic space. Performance is shown relative to the performance of a GFP control with the same compositional context arrangement of nodes (dashed green line). All 2X inverters are run with an extended equilibration (see Methods).

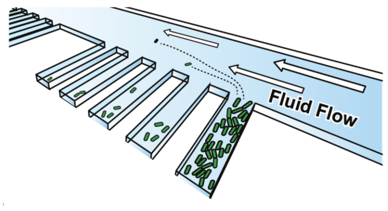


Supplementary Figure S8: **Mother machine.** Single-cell microfluidic study of circuit expression dynamics is accomplished using a device manufactured in-house at the CNF (Cornell Nanofabrication Facility). Single cells and their progeny are trapped in chambers perpendicular to the main channel. Side-by-side performance of cells under induction is included in Movie S1.

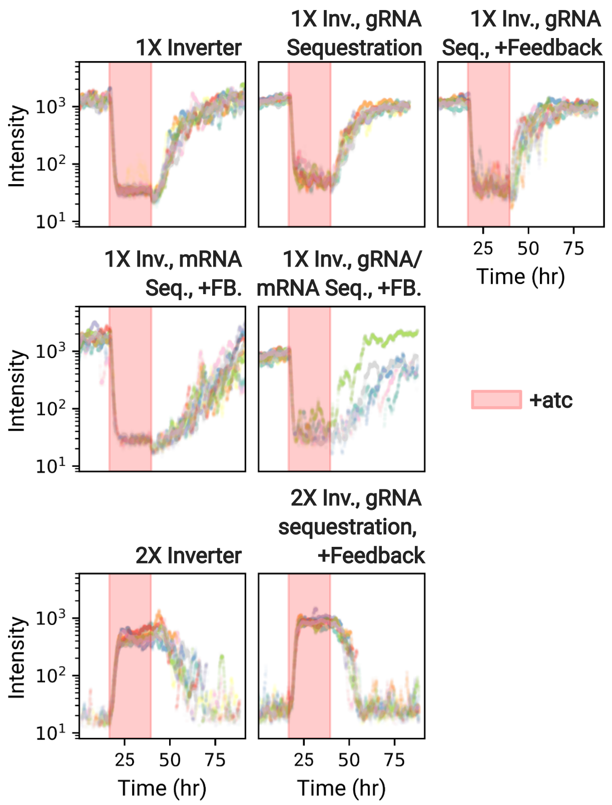


Supplementary Figure S9: **Channel median single-cell intensity.** Traces depicting channel median intensity of GFP expression across the entire course of the dynamic induction experiment. Reported data is the product of growth in 9 channels monitored over the course of the experiment. Time begins 1000 minutes before atc induction. Traces in which fewer than 10 cells are detected in a chamber are excluded and not shown.

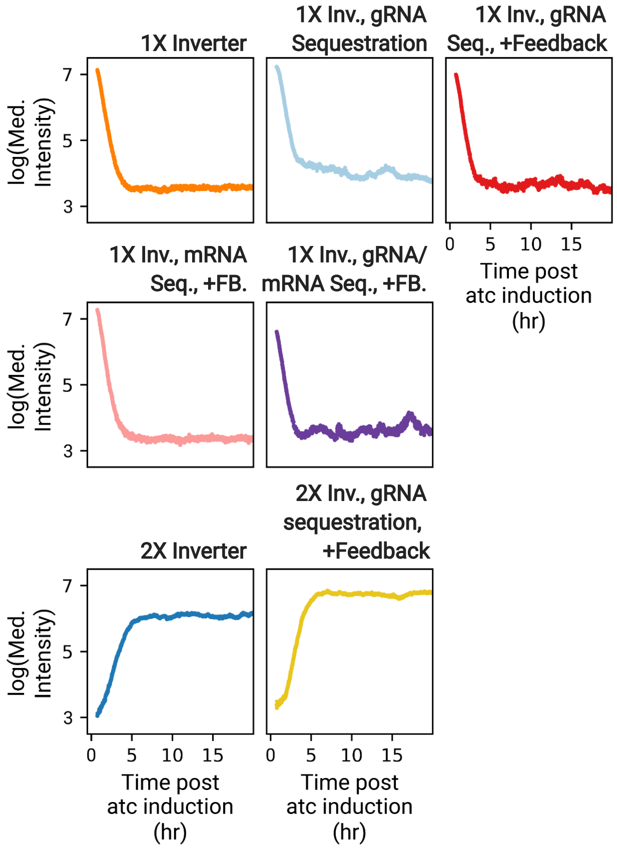


Supplementary Figure S10: **Single cell response to atc induction.** $t_{\frac{1}{2}}$ is calculated using the depicted spline curve fit to the log of the median intensity of single cell fluorescent intensity. Data points are median intensity of single cells across all microfluidic channels. The first 15 minutes of data post-induction are excluded from the spline fit because of transient effects from when the media is changed.

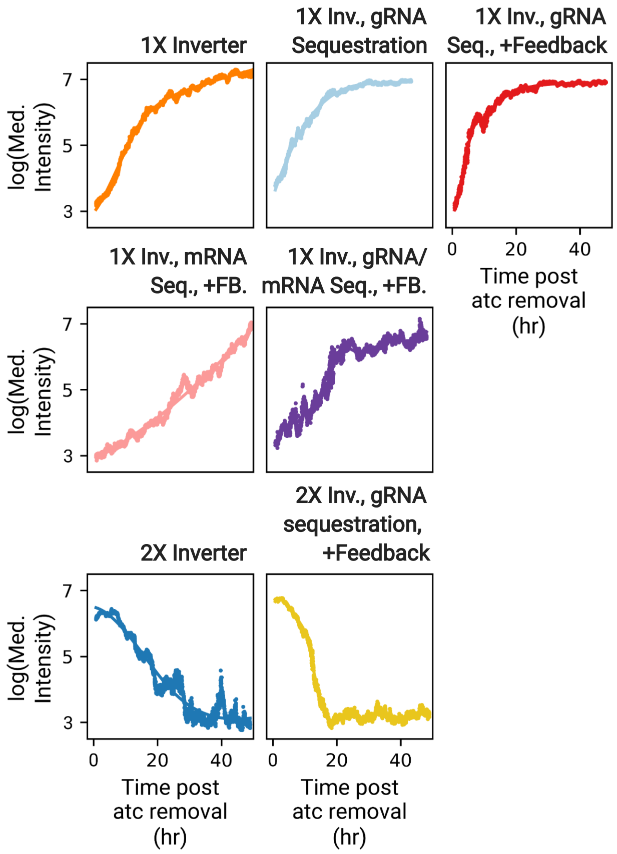


Supplementary Figure S11: **Single cell response to atc removal.** $t_{1/2}$ is calculated using the depicted spline curve fit to the log of the median intensity of single cell fluorescent intensity. Data points are median intensity of single cells across all microfluidic channels. The first 15 minutes of data post-induction are excluded from the spline fit because of transient effects from when the media is changed.

### 3 Supplemental Movie

#### 3.1 Side-by-side comparison of circuit performance under induction.

Five representative chambers for each construct illustrate single-cell response to atc addition and removal. We typically observe the circuits in its non-induced state overnight, then add atc to induce the circuit for 23 hours $\pm 15$ min and finally record the recovery after removal of atc for 40 to 60 hours, depending on the apparent time to equilibration. 20 to 40 chambers are monitored in parallel in a given experiment.
